# Elucidating essential genes in plant-associated *Pseudomonas protegens* Pf-5 using transposon insertion sequencing

**DOI:** 10.1101/2020.07.16.205906

**Authors:** Belinda K Fabian, Christie Foster, Amy J Asher, Liam DH Elbourne, Amy K Cain, Karl A Hassan, Sasha G Tetu, Ian T Paulsen

**Author notes:** Address correspondence to Ian Paulsen, or Sasha Tetu,.

## Abstract

Gene essentiality studies have been performed on numerous bacterial pathogens, but essential gene sets have been determined for only a few plant-associated bacteria. *Pseudomonas protegens* Pf-5 is a plant-commensal, biocontrol bacteria that can control disease-causing pathogens on a wide range of crops. Work on Pf-5 has mostly focused on secondary metabolism and biocontrol genes, but genome-wide approaches such as high-throughput transposon mutagenesis have not yet been used in this species. Here we generated a dense *P. protegens* Pf-5 transposon mutant library and used transposon-directed insertion site sequencing (TraDIS) to identify 446 genes essential for growth on rich media. Genes required for fundamental cellular machinery were enriched in the essential gene set, while genes related to nutrient biosynthesis, stress responses and transport were under-represented. Comparison of the essential gene sets of Pf-5 and *P. aeruginosa* PA14, an opportunistic human pathogen, provides insight into the biological processes important for their different lifestyles. Key differences include cytochrome *c* biogenesis, formation of periplasmic disulfide bonds, lipid biosynthesis, ribonuclease activity, lipopolysaccharides and cell surface structures. Comparison of the Pf-5 *in silico* predicted and *in vitro* determined essential gene sets highlighted the essential cellular functions that are over- and underestimated by each method. Expanding essentiality studies into bacteria with a range of lifestyles can improve our understanding of the biological processes important for survival and growth in different environmental niches.

**Importance:** Essential genes are those crucial for survival or normal growth rates in an organism. Essential gene sets have been identified in numerous bacterial pathogens, but only a few plant-associated bacteria. Employing genome-wide approaches, such as transposon insertion sequencing, allows for the concurrent analysis of all genes of a bacterial species and rapid determination of essential gene sets. We have used transposon insertion sequencing to systematically analyze thousands of *Pseudomonas protegens* Pf-5 genes and gain insights into gene functions and interactions that are not readily available using traditional methods. Comparing Pf-5 essential genes with those of *P. aeruginosa* PA14, an opportunistic human pathogen, provides insight into differences in gene essentiality which may be linked to their different lifestyles.

## Introduction

*Pseudomonas* is a ubiquitous and extremely diverse genus with species occupying a range of niches and lifestyles, spanning from opportunistic human pathogens, such as *P. aeruginosa*, through to plant growth promoting strains, such as *P. protegens* Pf-5 (1). Pseudomonads have a core set of common genes (2), but the mechanisms that underly their niche specializations are not well understood (3). Determining the essential gene complement of related microbes residing in different niches has the potential to shed light on differences in the biochemical functions required for survival and growth in specific environments. As observed in other bacterial taxa, each pseudomonad is expected to have a number of common essential genes together with an additional set of strain-specific essential genes, which may reflect differences in lifestyle (4, 5).

Essential genes are those crucial for survival or normal growth rates in an organism (6, 7). When bacterial genes are manipulated, essential genes are considered to be those for which mutations cannot be made as deletion of these genes will be lethal (8). Experimentally determining which bacterial genes are essential is important for understanding the mechanisms that control bacterial growth, identifying the mechanisms by which microbes specialize for their environmental niches, and can assist with validating computational models of gene essentiality (7, 9, 10). While gene essentiality studies have been performed on numerous bacterial pathogens, essential gene sets have been determined for only a few plant-associated bacteria, including *Herbaspirillum seropedicae* SmR1, a plant growth promoting endophyte, and three nitrogen-fixing root endosymbionts (11, 12).

*P. protegens* Pf-5 (hereafter referred to as Pf-5) is a plant-commensal, biocontrol bacteria originally isolated from the roots of cotton plants (13). Pf-5 is known to produce a range of secondary metabolites with antibacterial and antifungal activities (14) and can control disease-causing pathogens on a wide range of crops, including cotton, wheat, cucumber and tomatoes (15–18). Analysis of the Pf-5 genome has shed light on many potential functions and molecular systems utilized in its rhizospheric lifestyle (14, 19). Work on specific genes and gene networks has indicated functions for a number of Pf-5 genes, particularly regulatory and metabolic genes involved in the production of secondary metabolites. However, genome-wide approaches such as high-throughput transposon mutagenesis have not yet been used to elucidate gene function or essentiality in this important organism.

Here, we define the essential genome of *P. protegens* Pf-5 using transposon-directed insertion site sequencing (TraDIS). This method combines high-density random transposon insertion mutagenesis with high-throughput sequencing to concurrently link genotype and phenotype for thousands of genes (20, 21). TraDIS and other transposon mutagenesis techniques have successfully been used to determine the essential gene sets of a wide range of bacteria (22). We also compare the Pf-5 essential gene set with that of *P. aeruginosa* PA14 (hereafter referred to as PA14), an opportunistic human pathogen, providing insight into biological processes critical for their different lifestyles.

## Materials and Methods

### Transposon mutant library generation and sequencing

*Pseudomonas protegens* Pf-5 was isolated from soil of a cotton field in Texas, USA (13) and a complete genome sequence has been generated for this organism (19; ENA accession number CP000076). Construction of a *P. protegens* Pf-5 dense Tn*5* mutant library was carried out using the technique previously described (20, 21). Briefly, a custom transposome was constructed using EZ-Tn*5* transposase (EpiCentre) and a transposon carrying a kanamycin (Km) resistance cassette isolated from the plasmid pUT_Km (23). The custom transposome was electroporated into freshly prepared electrocompetent Pf-5 cells, and the cells were plated on LB-Km agar (16 ug mL^-1^). Approximately 125,000 colonies were collected from each of four independent batches, combined and stored as glycerol stocks at −80°C. Genomic DNA was isolated from two aliquots of stock containing approximately 2.8 × 10^9^ cells and the transposon insertion sites were sequenced using the methods described previously (21) on an Illumina MiSeq platform to obtain 52 bp single-end genomic DNA reads.

### Bioinformatic analysis

The transposon insertion sites were mapped to the Pf-5 genome and statistically analyzed using the Bio-Tradis pipeline (21). A 1 bp mismatch in the transposon tag was allowed and insertions in the extreme 3’ end (final 10%) of each gene were discounted as they may not inactivate the gene. Reads with more than one mapping location were mapped to a random matching location to avoid repetitive elements artificially appearing to be essential (0.4% of reads; 21). The pipeline calculates an insertion index value for each gene. This is the number of transposon insertion sites per gene normalized by the length of that gene. A linear regression of the gene insertion indexes of the replicates was completed in R (24). There was a correlation co-efficient of R^2^ = 0.88 (*p* < 2.2 × 10^-16^) between the insertion indexes of the replicates which validates the reproducibility of our replicates and is consistent with the high reproducibility between independent replicates in transposon insertion sequencing studies (25). Essential genes are those with an insertion index lower than the essentiality cut-off value determined by the Bio-Tradis pipeline (26). In this study we required essential genes to have an insertion index lower than the cut-off value in both replicates.

### Essential gene analysis

A Cluster of Orthologous Groups (COG) code (27) and Kyoto Encyclopedia of Genes and Genomes (KEGG) Orthology (KO) term (28, 29) for each Pf-5 gene was gathered using eggNOG mapper v1 (30, 31; Dataset S1). We used eggNOG mapper v2 (31, 32) to find this information for 18 essential genes where there were no orthologs identified by eggNOG mapper v1. COG functional category enrichment analysis of the essential gene set compared to the whole gene set was conducted using Fisher’s exact test (*p* < 0.05) and corrected for multiple testing using a 5% false-discovery rate (FDR; 33). The sum of all categories does not equal the total number of genes in the genome as some genes are assigned multiple COG codes. The KEGG mapper tool (34) was used to map the essential genes to KEGG pathways.

The presence and type of signal peptides in essential genes were identified using SignalP v5.0 (Dataset S1; 35) and the presence of transmembrane domains in proteins was determined using TMHMM v2.0 (Dataset S1; 36). As a transmembrane helix close to the N terminus is likely to be a signal peptide, proteins were only classed as membrane proteins if transmembrane helices were detected outside the first 50 residues. Enrichment for genes with signal peptides or transmembrane helices in the essential gene set compared to the whole genome was tested using Fisher’s exact test (*p* < 0.05) with Bonferroni correction for testing multiple values.

Pf-5 essential genes were also compared with PA14 essential genes determined by Poulsen and colleagues, which were based on growth in LB media (5). The Poulsen study determined the essential protein-coding genes using two statistical methods: the family-wise error rate (FWER) method, which identified 437 genes as essential, and the false discovery rate (FDR) method, which identified an additional 159 genes as essential (total of 596 essential genes). Orthologs of Pf-5 essential genes in PA14 were determined using Proteinortho v5 (37) and these were used to compare Pf-5 essential genes to the sets from PA14 derived from each statistical method.

*In silico* prediction of Pf-5 essential genes was conducted using Geptop 2.0 (http://cefg.uestc.cn/geptop/) with amino acid sequences and an essentiality score cutoff of 0.24 (38). The predicted Pf-5 essential genes were compared with the *in vitro* determined Pf-5 essential genes. The gene PFL_0842 was originally annotated as a pseudogene, so it was not assessed by Geptop (marked ‘N/A’ in Dataset S1).

### Data availability

All sequence data generated in this study have been submitted to the EBI European Nucleotide Archive (https://www.ebi.ac.uk/ena/) under the project accession number PRJEB39292, within which are the two samples analyzed here ERR4327923 and ERR4327924.

## Results and Discussion

### Identification of *P. protegens* Pf-5 essential genes

Our dense mutant library of *P. protegens* Pf-5 was generated using random saturation mutagenesis with a Tn*5* transposon containing a kanamycin resistance cassette. Over 500,000 transposon mutants were pooled in the construction of the library and the analysis of sequencing data showed there are ~256,000 unique transposon insertion sites in the mutant library (Table 1). Transposon insertions occurred evenly throughout the genome (Figure 1a), with an average of one transposon insertion every ~27 bp and an average of 45 transposon insertion sites in each non-essential protein-coding gene. Multiple insertion sites in a gene are independent evidence that a gene is not essential under the specific conditions used (39, 40). Rarefaction analysis showed that the sequencing reached saturation in terms of the number of unique transposon sites identified in the library (Figure 1c).

**Figure 1.**
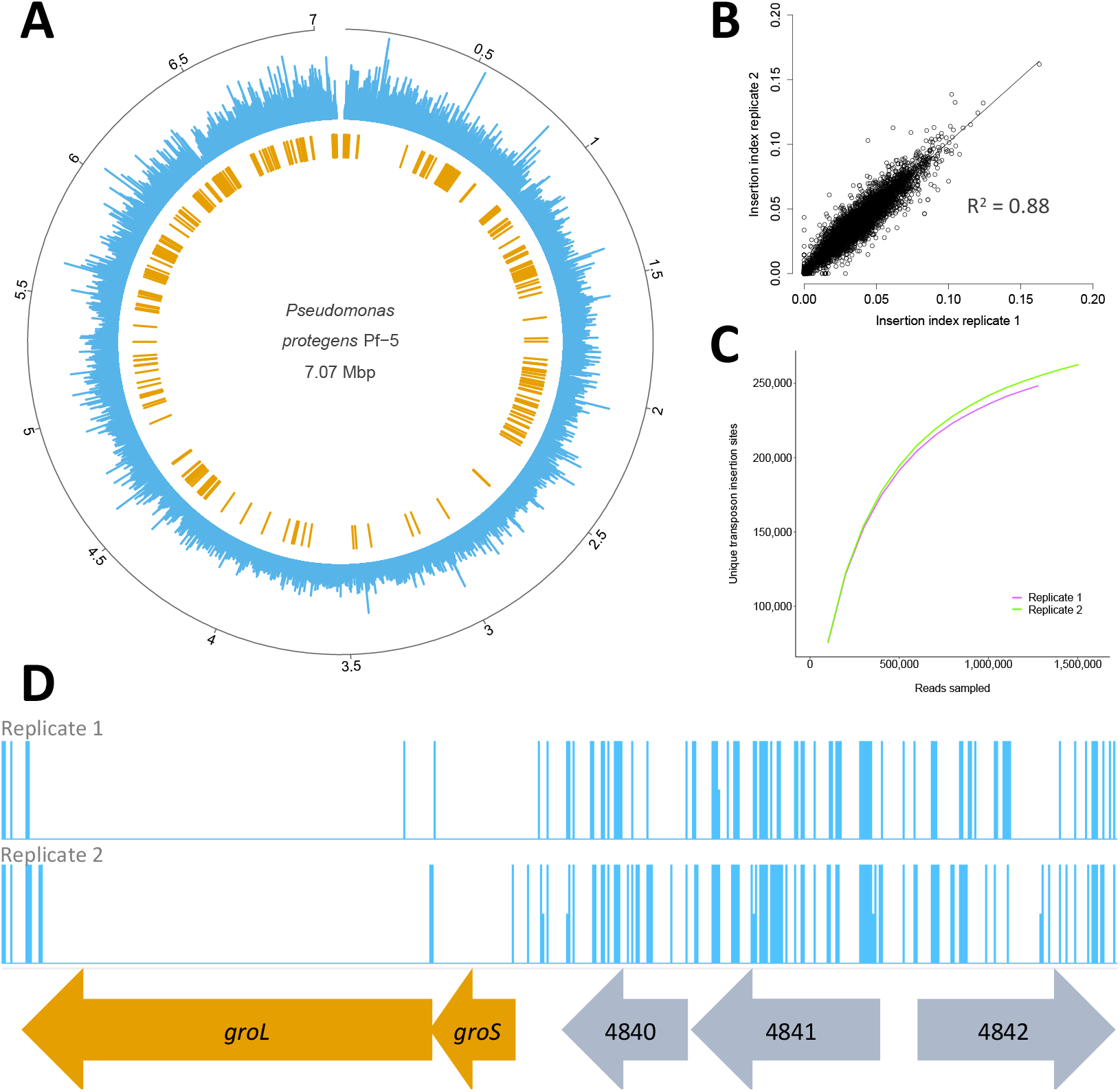
*P. protegens* Pf-5 transposon mutant library overview. (A) Distribution of transposon insertion sites across the genome. The outermost circle in grey shows the Pf-5 genome in Mbp. The length of the blue bars in the middle circle represents the number of sequencing reads at each unique transposon insertion location. The inner circle in orange shows the locations of genes characterized as essential with TraDIS. Circular figure created using the R package circlize (41). (B) Correlation of gene insertion indexes for the two replicates of the library. Insertion index is calculated as the number of transposon insertion sites in a gene divided by the gene length. A line through the origin with a slope of 1 is also shown. (C) Rarefaction analysis showing the relationship between sequencing depth and the number of insertion sites in the transposon mutant library. Analysis conducted using the seq_saturation_test.py script available at https://github.com/francesca-short/tradis_scripts (42) and visualized using the R package ggplot2 (43). (D) A section of the Pf-5 genome containing both essential (orange) and non-essential (grey) genes. The frequency of sequence reads at each transposon insertion site is capped at 1 and genes are annotated with Pf-5 gene names or locus tag numbers. The genes *groL* and *groS* have very few or no insertions and are essential genes. The genes PFL_4840, PFL_4841 and PFL_4842 have a high transposon insertion density and are therefore classed as non-essential genes. Transposon insertion sites are visualized using Artemis (44).

**Table 1.** *Pseudomonas protegens* Pf-5 transposon mutant library metrics based on Bio-Tradis analysis.

| Replicate | No. of reads with matching transposon tag | No. (%) of aligned reads | No. of unique insertion sites | Insertion index for essentiality cut-off |
| --- | --- | --- | --- | --- |
| 1 | 1,320,631 | 1,278,951 (96.8%) | 242,827 | 0.0076 |
| 2 | 1,555,998 | 1,504,986 (96.7%) | 256,020 | 0.0089 |

We found 446 out of 6,109 coding sequences (7.3%) were critical for Pf-5 survival and growth on LB agar (Dataset S1). Other pseudomonads that have their essential gene sets characterized also have similar proportions of their genomes determined to be essential genes. For example, *P. aeruginosa* PA14 has 596 out of 5894 protein-coding genes (10%) identified as essential when grown on LB (5) and 6% of the *P. aeruginosa* PAO1 genome was found to be essential for growth on a complex medium (45). The few plant-associated bacteria that have their essential gene sets identified have similar proportions of essential genes, for example *H. seropedicae* SmR1, a plant endophyte, required 372 out of 4804 protein-coding genes (7.7%) for survival and growth on rich media (12) and 5.6% of the genome of the root endophyte *Rhizobium leguminosarum* bv. *viciae* 3841 was determined to be essential (11). Pf-5 and other plant-associated bacteria often have large genomes with numerous stress response and biosynthetic pathways that are important for survival in their highly variable environmental niches (19, 46, 47). Many of these pathways were likely not critical for Pf-5 survival in this study due to the nutrient-rich and low stress conditions.

Transposon insertion density was slightly higher for genes located near the origin of replication compared to genes located closer to the terminus of replication (Figure 1a). This is regularly observed in genomic and transcriptomic studies due to ongoing DNA replication in the bacterial population. In keeping with this, the density of essential genes was also somewhat lower towards the replication terminus (Figure 1a). Lower numbers of essential genes in terminus regions may also reflect common patterns of genome arrangement; for example, terminus regions of bacterial genomes are often poorly conserved and experience higher rates of lateral gene transfer and rearrangement (48, 49).

The set of genes determined to be essential under the test conditions included PFL_0842, previously annotated as a pseudogene. Analysis showed that this sequence likely represents a functional gene, as it has positional orthologs with the same genomic context (part of an operon with *nusA* and *infB*) in numerous *Pseudomonas* species annotated as *rimP* ribosome maturation factor (50). Based on this, PFL_0842 has been annotated as *rimP* in this study.

### Functions of essential genes

A functional overview of the Pf-5 genome was obtained by classifying the genes using Clusters of Orthologous Groups (COG) categories (27). 87.3% of Pf-5 coding genes were able to be assigned a COG code. Several functional categories were significantly over-represented in the set of essential genes relative to the overall genome, most notably translation (J); cell cycle control, cell division and chromosome partitioning (D); coenzyme transport and metabolism (H); nucleotide transport and metabolism (F) as well as intracellular trafficking, secretion, and vesicular transport (U); cell wall/membrane/envelope biogenesis (M); and replication, recombination and repair (L; Figure 2). Other functional categories were significantly under-represented in the set of essential genes including secondary metabolite biosynthesis, transport and catabolism (Q); amino acid transport and metabolism (E); inorganic ion transport and metabolism (P); transcription (K); and signal transduction mechanisms (T; Figure 2). Genes with unknown function (S) make up a significantly lower proportion of the essential gene set (10.6%) compared to the whole Pf-5 genome (22%; Figure 2).

**Figure 2.**
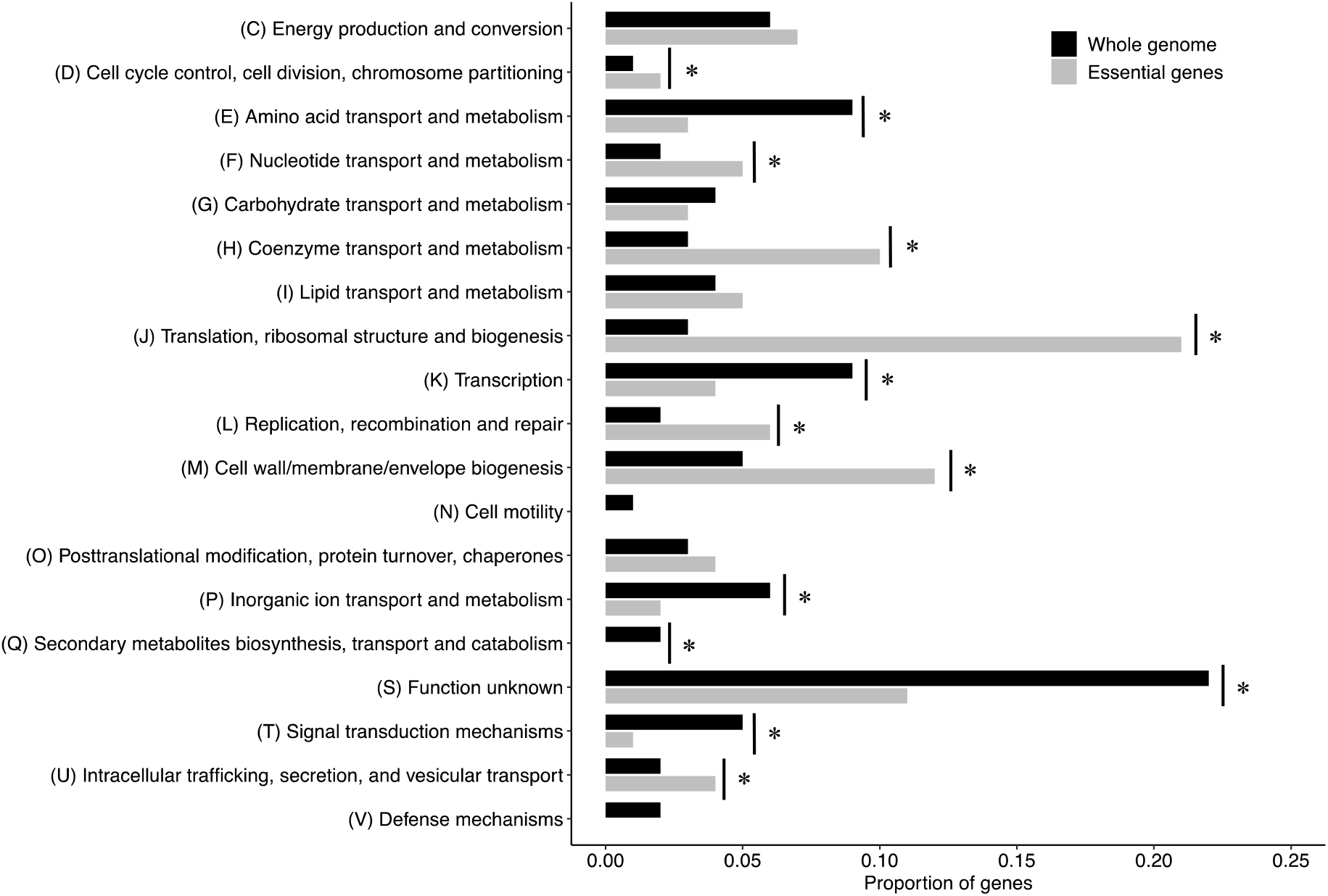
Cluster of Orthologous Groups (COG) functional enrichment analysis of *P. protegens* Pf-5 essential genes compared to the whole genome. COG category enrichment was tested using Fisher’s exact test and corrected for multiple testing using a 5% false-discovery rate; * indicates significant enrichment or depletion with *p* < 0.05; six COG categories are not included in the figure: categories A and B contain 2 and 5 non-essential genes, respectively, and categories R, W, Y and Z contain no genes.

The pattern of enrichment in essential genes belonging to COG categories for fundamental cellular machinery, such as translation and ribosome structure, coenzyme transport and metabolism and cell envelope biogenesis, reflects the fact that these functions are critical for survival and growth. In contrast, genes belonging to the COG category of transcription were notably under-represented in the essential gene set, presumably as transcriptional regulatory genes are typically required under changing environmental conditions (51), and therefore were not essential during growth in stable laboratory conditions with rich media.

At the metabolic pathway level, most of the key pathways were essential in Pf-5, including the TCA cycle, fatty acid biosynthesis, gluconeogenesis, peptidoglycan biosynthesis and heme biosynthesis, with all or most of the genes within each of these pathways observed within the essential gene set. Genes required for glycolysis, however, were not essential in this study, presumably as the mutant library was grown on LB media which contains high concentrations of peptides and amino acids which feed into other metabolic pathways. Similarly, under-representation of genes required for amino acid and inorganic ion transport and metabolism was observed, likely due to these biosynthetic pathways not being required for growth on rich media; this has also been observed in other gene essentiality studies across a wide range of bacterial taxa (52). Secondary metabolite biosynthesis, transport and catabolism genes were also under-represented in the essential gene set, potentially due to the limitation of TraDIS methodology in the detection of genes related to public goods (53).

Although Pf-5 encodes 780 transporter proteins (54), only four transport systems were essential: the LptBFG, MsbA, LolCDE, and CcmABC complexes. The first three of these transport systems are involved in the biogenesis of the outer membrane, while CcmABC is a heme chaperone-release system required for the biogenesis of cytochrome *c* (55, 56). The non-essentiality of other transporters was likely due to functional redundancy, the presence of gene duplicates, or that they were superfluous for growth in rich media.

Among the set of essential genes, a number remain annotated as hypothetical genes (35 genes or 7% of the essential gene set). The essentiality of these hypothetical genes indicates they are associated with as yet unknown functions that are critical for Pf-5 survival and growth on rich media. Given this, characterization of these genes would be of particular interest and may provide further useful insights into fundamental biological processes in this plant-associated bacteria.

### Essential genes associated with mobile genetic elements

The Pf-5 genome contains six prophage regions (Prophage 01-06) and two large genomic islands (PFGI-1 and -2), together comprising a total of 226 genes (57). Ten of the essential Pf-5 genes reside within these genome regions (Dataset S1). This includes PFL_4679, encoding type IV pilus biogenesis protein PilR which is located within ICE-type genomic island PFGI-1. This gene is part of a *pil* cluster, orthologs of which in PA14 are also located within a genomic island and have been found to be involved with conjugative transfer of the excised island to recipients (58, 59), but were not found to be essential in PA14 Tn-Seq experiments using rich medium (5). As *pilR* was the sole essential gene from this cluster in Pf-5, it may be that mutants of this gene were not viable due to accumulation of a toxic intermediate product such as non-polymerized pilin monomers, rather than due to impaired pilus formation capacity, which would presumably select against mutation of genes for the other pilus components.

The F-pyocin-like Prophage 01 contains the essential gene PFL_1210 which encodes the transcriptional regulator PrtR (57), a Cro/CI-like repressor of pyocin production, which has been previously found to be essential for *P. aeruginosa* PAO1 in rich media (45, 60). Essentiality of the *prtR* gene was presumably important for control of production of these polypeptide toxins to prevent self-lysis. Other prophage regions contain additional regulators observed to be essential, including two more Cro/C1-type transcriptional repressors (PFL_2126 in Prophage 04 and PFL_3780 in Prophage 06) and a peptidase S24-like protein/LexA-like repressor (PFL_1986 in Prophage 03; 57), which may similarly be required to prevent production of toxins or phage lytic conversion.

Of the remaining five MGE-associated essential genes, two have only predicted functions: the putative ATP-dependent nuclease PFL_1842 in Prophage 02 and the putative nuclease PFL_4984 within PFGI-2 (57); the remainder are conserved hypothetical proteins and potentially important targets for future characterization efforts. The presence of multiple essential genes within these MGE regions shows the capacity for such horizontally inherited material to acquire critical functions within bacterial cells, either in aiding stability of the MGE within the genome (for example, toxin/antitoxin systems) or contributing other conditionally essential functions.

### Protein subcellular localization of essential genes

Enrichment analysis for genes containing signal peptide sequences showed that genes that encode proteins containing lipoprotein and general secretion (sec) signal peptides were significantly under-represented in the essential gene set when compared to the Pf-5 genome as a whole (Table 2). Similarly, genes encoding proteins with transmembrane helices were significantly under-represented in the Pf-5 essential gene set (Table 2). This under-representation in the essential gene set is consistent with the subcellular localization of essential genes of 20 other gram-negative bacteria (61). As transmembrane proteins and those containing signal peptides are secreted outside cells or anchored in cell membranes they are often involved in interactions with other microorganisms and hosts (62) and therefore may not have been required under axenic conditions.

**Table 2.** Bioinformatic prediction of genes with signal peptide and transmembrane helices in the Pf-5 essential gene set and the whole Pf-5 genome.

| Protein subcellular localization | Number in essential gene set | Proportion of essential genes | Number in whole genome | Proportion of all genes |
| --- | --- | --- | --- | --- |
| SP | 19 | 4.3%<sup>†</sup> | 673 | 11.0% |
| TAT | 3 | 0.7% | 55 | 0.9% |
| LIPO | 8 | 1.8%<sup>†</sup> | 260 | 4.3% |
| **Total signal peptides** | **30** | **6.7%** | **988** | **16.2%** |
| Transmembrane proteins<sup>^</sup> | 40 | 9.0%<sup>†</sup> | 1089 | 17.8% |

SP = Sec signal peptide; TAT = Tat signal peptide; LIPO = Lipoprotein signal peptide
Enrichment of these genes in essential gene set was tested using Fisher's exact test with Bonferroni correction for testing multiple values.
<sup>†</sup> indicates significant underrepresentation in the essential gene set with $p < 0.05$
<sup>^</sup> one or more transmembrane helices detected outside the first 50 residues

### Comparison of essential genes in *P. protegens* Pf-5 and *P. aeruginosa* PA14

Based on our analysis, there were 446 essential protein-coding genes in Pf-5 when grown on LB media. A similar Tn-Seq study in PA14 has recently investigated essential protein-coding genes for this opportunistic human pathogen on LB media (5). This paper utilized two statistical methods to identify essential genes, the first based on the family-wise error rate (FWER) method which identified 437 genes as essential and the second using a false discovery rate (FDR) method which identified 596 genes as essential. We compared our set of Pf-5 essential genes with both sets from Poulsen et al. (5) and found the FDR set more closely aligned with our essential gene calls (Dataset S1). The FDR set also included genes expected to be essential, such as F0F1 ATPase, cell division and ribosomal protein genes, which were not identified by the family-wise error rate analysis (5).

Using the FDR set from Poulsen et al. (5) we investigated the similarities and differences between the essential gene sets of Pf-5 and PA14, finding that most (80%) of Pf-5 essential genes overlap with those of PA14, while each species also had a number of unique essential genes (Figure 3). The majority of the 357 overlapping essential genes relate to basic cellular functions, such as translation, cell envelope biogenesis, co-enzyme transport and metabolism, energy production, replication and recombination, and lipid transport and metabolism.

**Figure 3.**
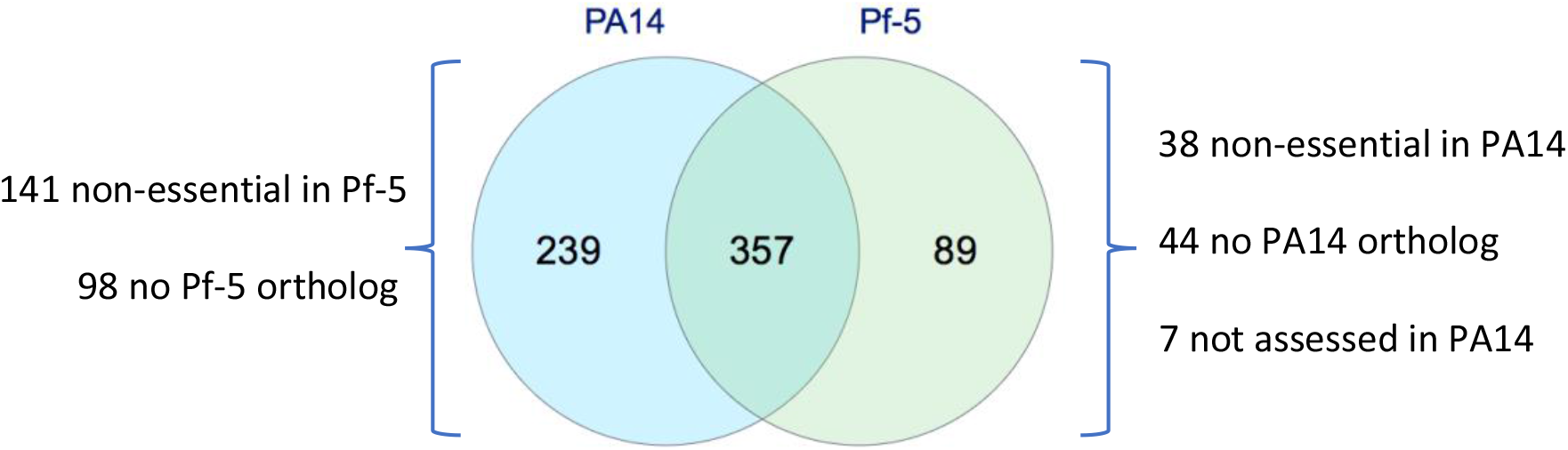
Comparison of the essential protein-coding genes of *P. protegens* Pf-5 with the essential genes of *P. aeruginosa* PA14 determined using a false discovery rate adjustment method (5). The PA14 orthologs of seven Pf-5 essential genes did not have their essentiality assessed by Poulsen et al. (5). Venn diagram created using Multiple List Comparator (www.molbiotools.com/listcompare.html).

#### Genes essential only in Pf-5

There were 38 genes within the Pf-5 essential set which were not flagged as essential in PA14 (Figure 3). This set included genes related to energy generation (cytochrome *c* biogenesis), formation of periplasmic disulfide bonds, three ribosomal proteins, amino acid biosynthesis, cell surface structures and hypothetical proteins.

Two-thirds of the genes in the cytochrome *c* maturation pathway are essential in Pf-5 (*ccmACEF*), but this pattern was not observed in PA14, where only one of these was in the essential set (*ccmB*). The *ccmABCDEF* genes encode proteins that carry out post-translational modifications on *c*-type cytochromes to facilitate their binding to heme (63). The essentiality of this set of genes in Pf-5 is consistent with the importance of generating cytochrome *c* to facilitate heme production.

Two of the genes essential only in Pf-5 encode proteins involved in the formation of disulfide bonds (*dsbD* and *trxB_1*). DsbD is a thiol:disulfide interchange protein which is the sole provider of periplasmic reducing power and has a role in cytochrome *c* maturation (64). In *Escherichia coli* DsbD is important, but not essential for cytochrome *c* maturation as developing cytochromes can use either the DsbD-independent or dependent pathways (64). The PA14 *dsbD* ortholog *dipZ2* and its paralog *dipZ* were both determined to be non-essential when grown on rich media (5). The essentiality of *dsbD* in Pf-5, but not in PA14, may reflect a different balance in the proportion of maturing *c*-type cytochromes that pass through each pathway. The Pf-5 gene *trxB_1* that encodes thioredoxin reductase was essential, but its paralog *trxB_2* was non-essential. In PA14 there are also two paralogs, *trxB_1* and *trxB_2*, but both of these genes were non-essential (5). *Trx* genes in bacteria supply reducing power to DsbD and play a role in stress responses, such as oxidative stress in *Pseudomonas syringae* pv. *tomato* (65). The difference in essentiality of the two Pf-5 *trxB* paralogs suggests that TrxB_1 may be the main provider of reducing power to DsbD and that TrxB_2 may be required under stress or other conditions.

Three ribosomal protein genes *rpmE_2, rpmF* and *rpmI* were essential in Pf-5, but not in PA14. In Pf-5 *rpmE_2* codes for a C+ form of the 50S ribosomal protein L31 which is stable when bound to a zinc ion; in contrast, *rpmE_1* encodes a C-form of the protein which lacks metal chelating capacity (66). The essentiality of *rpmE_2* in Pf-5 reflects the zinc replete conditions in this study, while *rpmE_1* has been found to be conditionally essential for Pf-5 in zinc limited conditions (66). In PA14, both of the genes annotated as *rpmE* were non-essential when grown on LB (PA14_66710 and PA14_17700; 5). The genes that encode 50S ribosomal proteins L32 (*rpmF*) and L35 (*rpmI*), whilst not essential in PA14 (5), have been found to be essential in a number of other bacteria. For example, *rpmF* was essential in *Burkholderia thailandensis* E264 (67) and both genes were essential in *Acinetobacter baylyi* ADP1 (68).

Genes encoding three proteins responsible for the formation of homoserine and its conversion to threonine (*thrBC* and *hom*) were essential in Pf-5, but not in PA14. These amino acids are important intermediates in amino acid biosynthesis and are precursors for the formation of methionine, serine, glycine and cysteine (69). These results suggest that PA14 was better able to take up threonine from the media in these conditions than Pf-5.

The gene *eda* which encodes 2-dehydro-3-deoxyphosphogluconate aldolase/4-hydroxy-2-oxoglutarate aldolase was essential in Pf-5 but its PA14 ortholog PA14_23090 was non-essential. Eda performs a key role in the catabolism of glucose and conversion into pyruvate through the Entner-Doudoroff pathway (70). This difference in essentiality is likely due to redundancy for this function in PA14; an *eda* paralog *kdgA* (PA14_23620) was also non-essential.

Three genes related to lipopolysaccharide formation (*kdsC, PFL_0526, PFL_0563*) were essential only in Pf-5. Similarly, in PA14 there are a number of essential genes that encode proteins important for lipopolysaccharide formation and other cell surface features that were not essential in Pf-5 (*htrB, omlA, wbpLV, rfbACD, pstB, wecB*). This difference in essential genes related to the cell membrane and cell surface likely reflects species-specific differences in many of these components, such as O-antigen, the polysaccharide component of lipopolysaccharide that extends from the surface of Gram-negative cells (71).

There were also differences in the essentiality of some conserved hypothetical proteins that are homologous in Pf-5 and PA14. The proteins encoded by PFL_1792 and PFL_2999 were essential in Pf-5, whereas their orthologs PA14_25620 and PA14_01220, respectively, were non-essential in PA14. Likewise, there were 16 essential conserved hypothetical proteins in PA14 but their orthologs were non-essential in Pf-5 (Dataset S1).

#### Pf-5 essential genes with no PA14 orthologs

There were 43 genes essential in Pf-5 that do not have homologs in PA14 (Figure 3), including genes related to cell membrane and surface structures, pyoluteorin biosynthesis, a TonB complex, and 25 conserved hypothetical proteins. The unique nature of the seven genes related to the cell surface and cell membrane (PFL_2356, PFL_5099-5103 and PFL_5030) is consistent with the species specificity of these components. The essential gene *pltL* from the pyoluteorin biosynthesis gene cluster has no PA14 homolog as this cluster is not present in PA14.

Genes that encode TonB complexes in both species were essential, but the genes that form some of the TonB systems are unique to Pf-5 and PA14. In Pf-5 the TonB1 complex (*tonB1, exbB1* and *exbD1*) was essential and there is no homologous TonB complex in PA14. In PA14 *tonB* (no Pf-5 homolog) and *exbD2* (homolog to PFL_2822) were essential. Many bacterial species only have one TonB system, but some species have multiple TonB systems with different functional specificities, for example *Vibrio cholerae* CA401S (72) and *P. aeruginosa* PAO1 (73). Both Pf-5 and PA14 possess multiple TonB complexes; Pf-5 has six annotated TonB complexes, while PA14 has two. This difference in essentiality of TonB systems of Pf-5 and PA14 is consistent with the differing functionalities of TonB systems observed in other species.

#### Genes essential only in PA14

There are 240 PA14 essential genes unique to PA14; only 141 of these have Pf-5 orthologs (Figure 3). This includes genes that encode proteins involved in four biosynthetic pathways (aromatic amino acids, biotin, lysine and lipids), cell division, homologous recombination, ribonuclease activity and 16 conserved hypothetical proteins.

Six genes in the aromatic amino acid biosynthetic pathway were essential in PA14, whereas all genes in this pathway were non-essential in Pf-5. This pathway, known as the shikimate pathway, produces chorismate which is the last common precursor of the aromatic amino acids tryptophan, tyrosine and phenylalanine as well as vitamins E and K and some siderophores (74, 75). The essentiality of *aroABCEK* and *pheA* show that chorismate is an important branch-point metabolite for PA14 survival and growth. In contrast, the non-essentiality of these genes in Pf-5 suggests that this pathway was not required under these experimental conditions and Pf-5 obtained aromatic amino acids from the media.

Genes encoding five proteins in the biotin biosynthesis pathway (*bioABCDF*) and its transcriptional repressor *(birA)* were essential in PA14, but non-essential in Pf-5. Biotin, also known as vitamin H, is a cofactor for enzymes involved in central metabolism carboxylation reactions (76). These results suggest that Pf-5 was better able to take up biotin from the media than PA14.

Four genes that encode proteins involved in cell division were essential in PA14, but non-essential in Pf-5 (*ftsK, minCD, mreD, PFL_0438*). Cell division proteins are essential for the normal replication and viability of bacterial cells; for example, *minCD* encodes proteins that ensure that cell division occurs in the middle of the cell not at a polar site (77) and *mreD* encodes a shape protein which is involved in the rod shape of cells. When *mreD* is knocked out cells take on a spherical form (78). Other rod-shaped bacteria, such as *A. baylyi* ADP1, have been able to dispense with these genes under lab conditions in essentiality studies (68) but *ftsK* and *mreD* have been reported to be essential in other studies in *P. aeruginosa* (45). The non-essentiality of these genes in Pf-5 suggests that cells without these proteins survived in the short timeframe of this experiment, albeit presumably with abnormal morphologies.

Five genes that code for lipid metabolism proteins were essential in PA14, but not Pf-5. The genes *fadA* and *fadB* encode the two peptides that form the fatty acid oxidation complex. This complex is part of the β-oxidation cycle which is responsible for the degradation of long chain fatty acids into acetyl-coenzyme A (79). In Pf-5 there is a single copy of *fadA*, but multiple copies of *fadB* (*fadB, fadB1x* and *fadB2x*). In *P. aeruginosa* PAO1 expression of each of the *fadAB* operon homologues is induced by the presence of different fatty acids (80, 81). There are also three acyl-CoA dehydrogenase family proteins (PFL_0245, PFL_2615 and PFL_5687) that were essential in PA14, but non-essential in Pf-5. The difference in the essentiality of these five genes in PA14 and Pf-5 suggests that the two species have different availabilities of fatty acids and maintain different balances in their lipid metabolism to achieve membrane homeostasis.

Genes *recBCD* and *ruvA* that encode homologous recombination enzymes were essential in PA14 but non-essential in Pf-5. A relatively small number of bacterial essential gene studies have identified these genes as essential (22), despite the importance of Rec-mediated repair of double stranded DNA breaks. The non-essentiality of *recBCD* and *ruvA* suggests this function is not essential for Pf-5 under laboratory conditions and timescales.

Genes that encode four ribonucleases were essential in PA14, but non-essential in Pf-5 (*rnc*, *rne*, *rnt* and PFL_3322). Ribonucleases (or RNases) have dual functions: they are involved in both the maturation and degradation of rRNAs, tRNAs, sRNAs and mRNAs (82). In *E. coli* there is evidence to suggest these ribonucleases may have overlapping functionalities (83), so these may have been non-essential in Pf-5 due to redundancy.

The genes *dapA_1* and *dapF_2* encoding proteins in the lysine biosynthetic pathway (84) were in the essential set in PA14 but not Pf-5. This difference in essentiality may occur as PA14 has a single copy of each gene, whereas Pf-5 has redundancy for these functions (two copies of each gene). A similar pattern is observed with the gene *fabF* which codes for beta-ketoacyl-acyl-carrier-protein synthase II, a part of the fatty acid biosynthesis pathway and a vital enzyme in the biogenesis of phospholipid membranes (85). The two copies of this gene in Pf-5, *fabF_1 and fabF_2*, were non-essential. Of the two PA14 copies of this gene *fabF1* was essential and *fabF2* was non-essential (5).

### Comparison of *in vitro* determined and *in silico* predicted essential genes

In parallel with the development of transposon library-based approaches for determining gene essentiality, computational tools have been developed to predict essential gene sets based on information such as gene orthology and phylogeny (38). Here we compared the set of essential genes identified *in vitro* by TraDIS with computational predictions by Geptop 2.0 (38). Geptop 2.0 identified 406 protein-coding genes as essential for growth and survival of Pf-5 (Dataset S1). When compared with the 446 essential protein-coding genes identified by TraDIS on rich media there were 308 genes in common, 138 genes only identified by TraDIS, and 98 genes that were only in the computationally predicted set (Figure 4). The 308 genes identified as essential by both methodologies include core cellular functions such as the processing of information (replication, translation and transcription), energy production, cell division and maintenance of the cell envelope.

**Figure 4.**
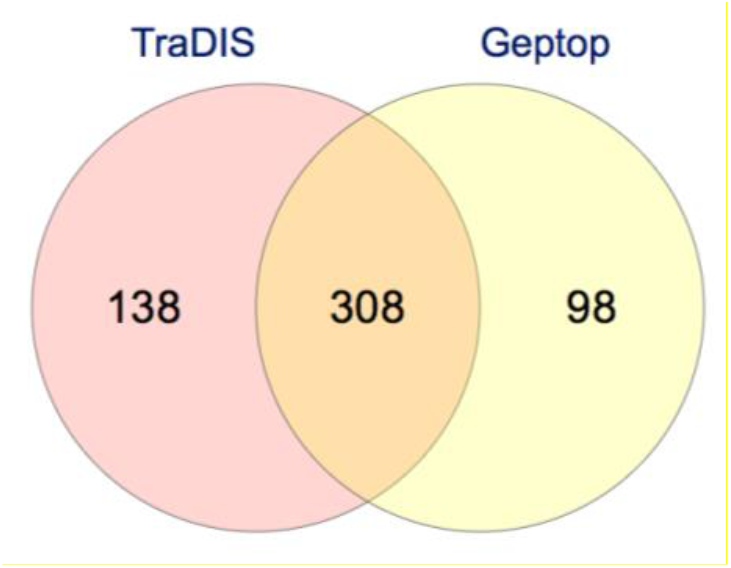
Comparison of *P. protegens* Pf-5 essential protein-coding genes identified *in vitro* by TraDIS and the essential genes predicted *in silico* by Geptop 2.0 (38). Venn diagram created using Multiple List Comparator (www.molbiotools.com/listcompare.html).

Out of the 138 essential genes identified only by TraDIS, 14 are related to lipopolysaccharide biosynthesis and cell surface structures, including *rfaCF, waaGP* and a range of lipoprotein, O-antigen and glycosyltransferase genes. The absence of these genes from the predicted essential gene set is consistent with the species-specific nature of these pathways and the use of evolutionary conservation in the Geptop computational approach (38). Thirty-two essential genes identified only by TraDIS are annotated as hypothetical proteins. As Geptop uses orthology and phylogeny to predict essential genes it is unsurprising that hypothetical genes were not included in the predicted essential gene set.

There are a number of genes our work indicates are essential that have not been identified by Geptop. For example, *fbp* encoding frustose-1,6-bisphosphatase, cell division genes (*zipA, ftsB* and *minE*), *fis* encoding the DNA binding protein Fis, cytochrome c biogenesis (*ccmABCEF*), genes encoding proteins involved in the maintenance of membrane stability (*pal* and *tolABQR*), sulfur relay genes (*tusABCDE*) and iron-sulfur cluster genes (*hscAB, iscA* and ferrodoxin genes *fdx* and PFL_5869). These genes have been found to be essential in other organisms such as *E. coli, Haemophilus influenzae* and *Shewanella oneidensis* (86–88). This suggests there may be functional categories of genes that are systematically missed by Geptop.

Ninety-eight Pf-5 protein-coding genes were identified as essential by Geptop but not found to be essential *in vitro* by TraDIS. There are three major reasons these genes were not identified as essential in the *in vitro* data. Firstly, there is some redundancy in the genome for essential functions. For example, the genes *ddlAB* encode D-alanine--D-alanine ligases which condense two molecules of D-alanine, an essential step in peptidoglycan biosynthesis (89, 90), yet neither gene was essential *in vitro*, presumably as loss of function of each single gene can be tolerated. Similarly, *map_1* and *map_2* both encode methionine aminopeptidases which perform the essential function of cleaving methionine residues from the N-terminal of nascent proteins (91), but loss of either of these genes did not preclude growth *in vitro*. Many of the genes encoding subunits of NADH-quinone oxidoreductase were non-essential in Pf-5 *in vitro* (*nuoBDGHIJKLMN*) but were predicted to be essential by Geptop. In addition to the *nuo* complex, which encodes a type I NADH dehydrogenase, there are two other NADH dehydrogenases encoded in the Pf-5 genome (92). These three NADH dehydrogenases provide Pf-5 with respiratory flexibility, such as in *P. aeruginosa* where two NADH dehydrogenases were redundant in aerobic conditions (93). As TraDIS library construction generates single knockout mutants there is limited capacity to identify essential genes where there is functional redundancy in the genome, which is a recognized limitation of this technique (94).

Secondly, some genes encode products with essential functions in bacterial cells, but these functions are only performed under certain conditions. For example, the highly conserved Clp proteolytic system degrades misfolded and abnormal proteins which accumulate in response to environmental stresses (95). This essential function is reflected in the Geptop prediction of *clpP* and *clpX* as essential, but these genes were not identified as essential by TraDIS, presumably as environmental stresses were low under our library growth conditions. This pattern is also observed with *dnaJ* which encodes a molecular chaperone which helps to ensure the correct folding of proteins, particularly under heat shock (96, 97), and the highly conserved gene *polA* encoding DNA Polymerase I which repairs damaged DNA during replication (98). These proteins perform important functions but were not essential under the stable conditions and short duration of this study. The media and growth conditions under which transposon libraries are created also influences the essentiality of genes (7). The rich media used in our *in vitro* experiments may have resulted in some genes that were predicted to be essential *in silico* not being identified as essential by TraDIS. For example, three amino acid biosynthetic genes (*aroBCK*) were predicted to be essential by Geptop, but were non-essential *in vitro*, likely due to the ability of Pf-5 to acquire amino acids from the media.

Lastly, the experimental timeframe is likely also a factor in some genes being identified by TraDIS as non-essential. These genes may have important roles at certain stages of cell or population growth and therefore are identified as essential based on the phylogenetic approach used by Geptop. For example, the genes *ftsHKX* and *parAB* perform important roles in cell division (99) and were predicted to be essential by Geptop, but they were determined to be non-essential by TraDIS. This indicates cells with these individual gene disruptions may undergo abnormal cell division, potentially resulting in aberrant morphologies, but the cells were not lost completely from the population. It is likely that such altered morphologies may become problematic in a longer-term study.

## Conclusion

In this study we created a saturated transposon mutant library and used TraDIS to successfully identify 446 genes that were essential for *P. protegens* Pf-5, a plant-associated bacteria, to survive and grow on rich media. The essential gene set showed enrichment of genes required for fundamental cellular machinery, which is consistent with the composition of essential gene sets in other bacteria. Genes related to nutrient biosynthesis, stress responses and transport were under-represented, potentially due to the specific growth conditions used in this study as well as functional redundancy within the genome.

We identified key differences between the essential gene sets of the plant-associated pseudomonad Pf-5 and the well-studied opportunistic pathogen PA14. These include genes related to energy generation (cytochrome *c* biogenesis), formation of periplasmic disulfide bonds, lipid biosynthesis, ribonuclease activity, lipopolysaccharides and cell surface structures. This information highlights differences in the processes required for survival and growth of pseudomonads that occupy different environmental niches.

Our comparison of the essential gene sets determined *in silico* and via the *in vitro* TraDIS approach shows that the prediction of essential genes by Geptop on the basis of conservation through evolutionary time overestimates the essentiality of some cellular functions and underestimates others. Despite this, there is still substantial overlap in the genes identified as essential by these two methods. While both techniques have recognized limitations, the information from TraDIS studies could be used to evaluate and improve *in silico* predictive models for essential genes.

Using TraDIS to systematically analyze thousands of genes provides insights into gene functions and interactions that are not readily available using traditional methods. The Pf-5 transposon mutant library will enable high-throughput studies in a range of growth conditions, such as competition with soil microbes or stress tolerance. Expanding essentiality studies beyond bacterial pathogens improves our understanding of the biological processes important for survival and growth in different environmental niches.

## Supporting information

Dataset S1 _P. protegens Pf-5 gene essentiality

## Acknowledgements

The TraDIS sequencing for this study was conducted at The Ramaciotti Centre for Genomics at the University of New South Wales. *Pseudomonas protegens* Pf-5 was obtained from Joyce Loper (Oregon State University). The authors thank Dr Francesca Short (Macquarie University) for advice on TraDIS data analysis and Professor Joyce Loper (Oregon State University) and Dr Virginia Stockwell (USDA) for helpful discussions. This work is supported by an Australian Research Council Discovery Grant (#DP160103746) to IP, ST and KH. IP is supported by an Australian Research Council Laureate Fellowship (#FL140100021), ST is supported by an Australian Research Council Discovery Early Career Researcher Award (#DE150100009) and BF is the recipient of an Australian Government Research Training Pathway Scholarship.

